# Discovery of novel members of the *Potyviridae* family reveals expanded diversity, a broad host range, and evidence of fungal and oomycete infections

**DOI:** 10.64898/2026.05.29.728796

**Authors:** Bernardo Rodamilans, Mario Rincón Barrado, Alberto Cobos Piñuela, Carmen Simón Mateo, Adrian A. Valli

## Abstract

The family *Potyviridae* represents the largest and most economically important group of plant-infecting RNA viruses. Despite extensive study of crop-associated members, the full diversity, host range, and evolutionary history of potyvirids remain poorly understood. Here, we conducted a large-scale mining of publicly available RNA-seq datasets to systematically search for novel potyvirid sequences. This approach enabled the identification and assembly of 47 previously undescribed members of the family, distributed across eight recognized genera and, importantly, two putative new genera. Beyond expanding the known genetic diversity of *Potyviridae*, our analyses revealed a distinct and deeply divergent lineage of potyvirid-like viruses associated with fungi and oomycetes, for which we propose the genus *Macrophovirus*. These viruses possess compact genomes and atypical genomic organizations, including the absence of canonical plant cell-to-cell movement factors and the presence of HCPro-like proteins arranged in tandem. Comparative structural and phylogenetic analyses indicate that these leader proteases are more closely related to fungal hypoviral counterparts than to canonical potyvirid HCPro factors. Together, our findings substantially expand the host range of *Potyviridae*, provide compelling evidence that potyvirid-like viruses likely infect fungi and oomycetes in nature, and offer new insights into the evolutionary pathways that have shaped this major virus family.

## INTRODUCTION

With 285 assigned members (204 with fully annotated genomes) classified into 13 genera according to the latest ICTV release (https://ictv.global/vmr, 03/07/2025), the *Potyviridae* family represents the largest and most economically significant group of plant-infecting RNA viruses. Members of this family cause substantial yield losses across a wide range of crops, including cereals, legumes, vegetables, and fruit trees, posing a serious threat to global agriculture and food security (Valli et al., 2021). Potyvirids (members of the *Potyviridae* family) possess monopartite genomes, with the exception of those in the *Bymovirus* genus, which are bipartite. Their genomes consist of positive-sense single-stranded RNA (+ssRNA) of approximately 10 kb in length (Revers and García, 2015, Yang et al., 2021). The genomic RNA is generally translated into a single large open reading frame (ORF), giving rise to a polyprotein that is subsequently cleaved by virus-encoded proteases into mature viral proteins. These typically include, from N- to C-terminus, the leader serine protease P1, the multifunctional cysteine-protease HC-Pro, the movement- and replication-associated P3, the membrane-associated factor 6K1, the helicase CI, the replication vesicle inducer 6K2, NIa (comprising the genome-linked protein VPg and the C4-like protease NIaPro), the RNA-dependent RNA polymerase (RdRP) NIb, and the capsid protein (CP) (Valli et al., 2021), with most sequence variation concentrated in the N-terminal region of the polyprotein (Pasin et al., 2022). The bipartite genome of bymoviruses consists of RNA1 and RNA2. RNA1 encodes the potyvirid P3- to-CP module, whereas RNA2 encodes the cysteine-protease P1 functionally equivalent to HCPro, and P2, whose function remains unknown (Valli et al., 2018, Niehl and Rabenstein, 2021, Chen et al., 2023). In addition to the main ORF, all potyvirids described so far encode at least one additional ORF, known as *pipo*, which is overprinted in the - 1/+2 reading frame of the P3 coding region. This ORF is preceded by an An or Un motif (where n ≥ 6), accessed via polymerase slippage, and expressed as a fusion with the N-terminal portion of P3, resulting in the production of P3N-PIPO (Valli et al., 2024). This protein, along with P3 and CI, participates in virus cell-to-cell movement via plasmodesmata (Chai et al., 2020).

Recently, a study on *Potyvirus rapae* (turnip mosaic virus, TuMV) highlighted the potential of the viral negative-strand RNA to encode small peptides with critical roles during infection (Gong et al., 2023). However, the presence of reverse ORFs does not appear to be a general property of all potyvirids, as recently reported (Garcia et al., 2025). Overall, as is typical of RNA viruses, potyvirids exhibit high genetic plasticity, which facilitates the emergence of novel strains and complicates disease management efforts. Large-scale reanalysis platforms, such as Serratus and VIRE (Edgar et al., 2022, Nishijima et al., 2026), have transformed virus discovery by enabling systematic mining of publicly available RNA-seq datasets for viral sequences at unprecedented scale. By screening millions of sequencing libraries, these platforms uncover vast numbers of previously unrecognized viruses across diverse environments and hosts, including organisms that are rarely targeted by traditional virological surveys. Similar high-throughput computational frameworks leverage advances in cloud computing, sequence alignment, and reference database curation to detect highly divergent viral genomes that would otherwise remain hidden. Together, these platforms are critical for improving our understanding of viral diversity and evolution, and for uncovering hidden viruses across diverse hosts and environments.

To gain insight into the diversity and host range of the family *Potyviridae*, we conducted a comprehensive survey of publicly available RNA-seq datasets via Serratus to identify novel potyvirid sequences. This approach led to the discovery of numerous previously unrecognized viruses spanning multiple genera within the family and, importantly, revealed a distinct group of potyvirid-like viruses associated with fungal samples. Collectively, these findings not only increase the known diversity of potyvirids and their plant hosts, but also provide compelling evidence for host range expansion and adaptation to fungi. The evolutionary and biological implications of these discoveries are discussed.

## MATERIALS AND METHODS

### Identification of new potyvirids from available RNA-seq data

The Serratus viral discovery platform (Edgar et al., 2022), and specifically its “Explorer RdRP Search” tool (https://serratus.io/explorer/rdrp), was employed to identify potential novel members of the *Potyviridae* family. A family-level search was performed using “Potyviridae” as the query, with default score (50-100) and identity (45%-100%) parameters. This configuration enabled the retrieval of RNA-seq data containing potyvirid replicase sequences. A table containing SRA IDs, their corresponding scores, the identity of detected potyvirid replicases relative to those in the Serratus RdRP database, and the number of reads, was downloaded from Serratus.

Two criteria were considered for candidate selection. First, putative novel potyvirid species are expected to exhibit replicase amino acid sequence identities below the species demarcation threshold of 89% established for members of the family *Potyviridae* (Adams et al., 2005b). Second, because individual SRA datasets may contain multiple matches against the Serratus RdRP reference set—which includes 11 diverse potyvirid RdRP representatives (Potyviridae-1 to Potyviridae-11)—only datasets in which all detected potyvirid RdRP hits displayed identities below the species threshold were retained. To ensure sufficient phylogenetic divergence from previously characterized viruses, a more stringent cutoff of <85% identity was applied. In addition, only datasets containing more than 500 reads mapping to the viral replicase coding region were selected, thereby reducing the likelihood of incomplete assemblies or incidental contamination with potyvirid-derived RNA.

Since Serratus relies on publicly available sequencing data deposited in NCBI before 2023, it was necessary to discard SRA hits including potyvirids annotated during 2023-2025 as they are no longer novel. To achieve this, all potyvirid *RdRP* sequences (rdrp.fasta) from each filtered SRA hit, which were retrieved from Serratus by using the SRA ID as query in SRA-level searches, were subjected to online BLASTX analysis (https://blast.ncbi.nlm.nih.gov/Blast.cgi). Only sequences displaying identity to other potyvirids below 85% were retained for further analyses.

Finally, selected SRA files were downloaded from NCBI, converted to FASTQ format, and used for transcriptome assembly with the Trinity RNA-Seq assembly tool (Grabherr et al., 2011) with default parameters. The resulting FASTA files containing assembled transcripts were screened to identify those corresponding to potyvirid genomes. For this purpose, a local TBLASTN search (Altschul et al., 1990) was performed using each transcriptome as a database and the replicase amino acid sequence of TuMV (NCBI accession: NP_734221) as the query. Similarly, the P1 amino acid sequence of *Bymovirus hordeiluteum* (NCBI accession: NP_734081) was used as query to identify the genomic RNA2 in the case of bymoviruses.

### Sequence alignments and phylogenetic analyses

GenBank accession numbers corresponding to the sequences used for multiple sequence alignments and phylogenetic tree construction are listed in Supplementary Table 1. Multiple sequence alignments were generated using Clustal Omega implemented in DNASTAR Lasergene MegAlign Pro (DNASTAR Inc., WI, USA). Maximum-likelihood phylogenetic inference was performed with RAxML. Circular unrooted trees were visualized using iTOL (Letunic and Bork, 2024).

### Protein 3D modelling

Protein models were obtained with AlphaFold3 (Abramson et al., 2024) using the protease domains of PVY HCPro, PvlaPV1 HCPro2 and CHV1 HCPro (see Supplementary Table 1 for NCBI accession numbers). The AlphaFold3 model of PvlaPV1 HCPro2 was used as the query in Foldseek (van Kempen et al., 2024) with default parameters to identify structurally similar proteins. Three-dimensional visualizations were produced in PyMOL (The PyMOL Molecular Graphics System, Version 3.0 Schrödinger, LLC).

## RESULTS

### Discovery of new members in the *Potyviridae* family

At the time of writing this manuscript, the complete genomes of 204 accepted potyvirids have been annotated according to the latest ICTV release (https://ictv.global/vmr, July 3, 2025). Of these, 162 belong to the *Potyvirus* genus, while the remaining viruses are distributed across 12 other genera: *Arepavirus*, *Bevemovirus*, *Brambyvirus*, *Bymovirus*, *Celavirus*, *Ipomovirus*, *Macluravirus*, *Phragmivirus*, *Poacevirus*, *Roymovirus*, *Rymovirus*, and *Tritimovirus*. This distribution highlights the remarkable evolutionary success of the *Potyvirus* genus.

To advance our understanding of the *Potyviridae* family, and particularly to expand knowledge of those underrepresented genera, we sought to identify novel members within this group. To this end, we leveraged the Serratus platform to screen for potyvirid genomes across more than five million publicly available RNA-seq datasets (see Materials and Methods for details). This large-scale mining effort led to the identification of 47 previously undescribed members of the family, distributed across eight recognized genera. A comprehensive summary of these findings is presented in Table 1. Although the overrepresentation of members of the genus *Potyvirus* within the family *Potyviridae* is again clearly evident, with 22 newly identified viruses assigned to this genus, the remaining 25 viruses do not cluster within *Potyvirus*. Moreover, a subset of these newly identified viruses appears to represent two putative novel genera within the family (see below for further details).

**Table 1:**
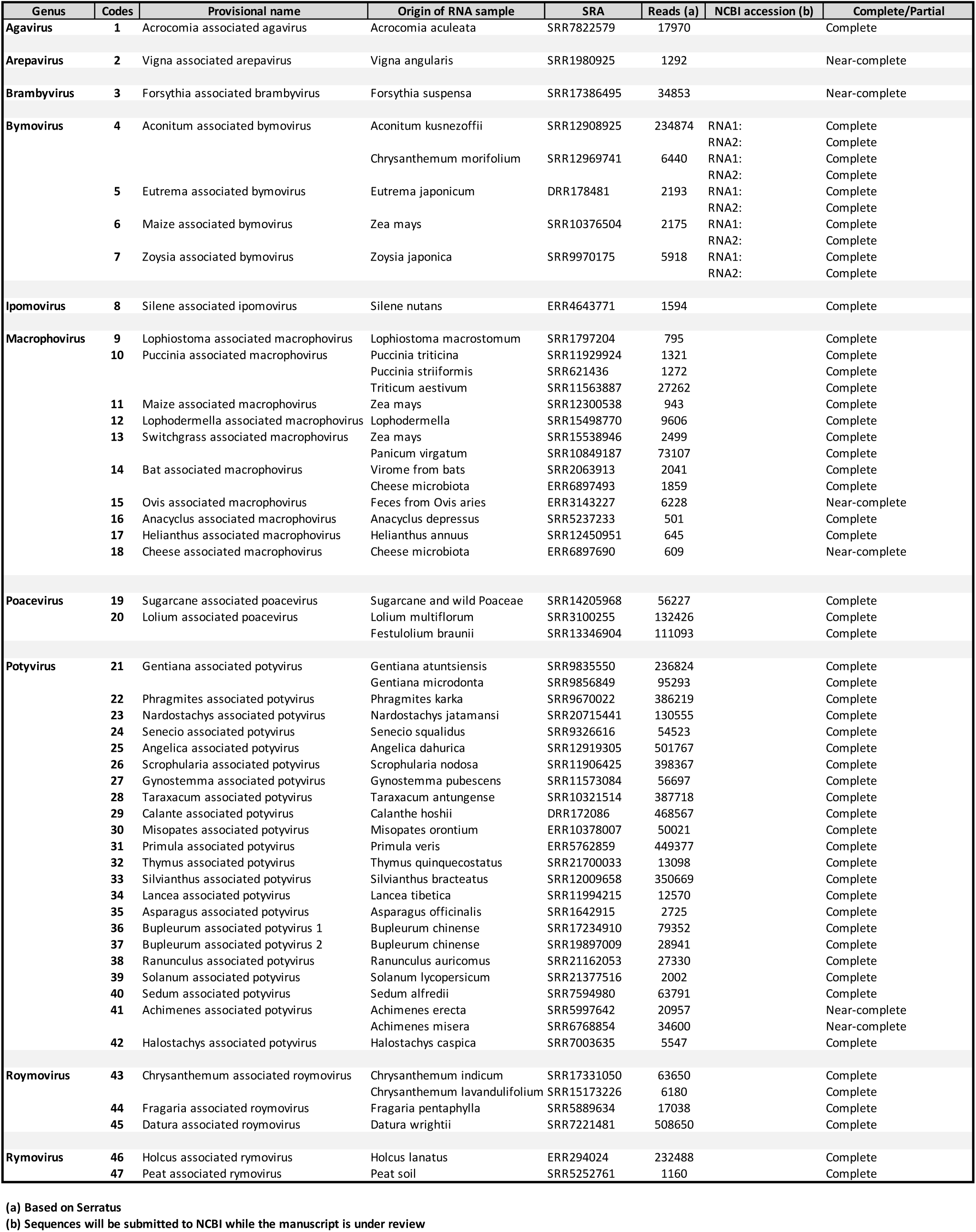
list of new potyvirids identified with the pipeline described in the materials and methods section.

| Genus | Codes | Provisional name | Origin of RNA sample | SRA | Reads (a) | NCBI accession (b) | Complete/Partial |
| --- | --- | --- | --- | --- | --- | --- | --- |
| **Agavirus** | 1 | Acrocomia associated agavirus | Acrocomia aculeata | SRR7822579 | 17970 |  | Complete |
| **Arepavirus** | 2 | Vigna associated arepavirus | Vigna angularis | SRR1980925 | 1292 |  | Near-complete |
| **Brambyvirus** | 3 | Forsythia associated brambyvirus | Forsythia suspensa | SRR17386495 | 34853 |  | Near-complete |
| **Bymovirus** | 4 | Aconitum associated bymovirus | Aconitum kusnezoffii | SRR12908925 | 234874 | RNA1: | Complete |
|  |  |  |  |  |  | RNA2: | Complete |
|  |  |  | Chrysanthemum morifolium | SRR12969741 | 6440 | RNA1: | Complete |
|  |  |  |  |  |  | RNA2: | Complete |
|  | 5 | Eutrema associated bymovirus | Eutrema japonicum | DRR178481 | 2193 | RNA1: | Complete |
|  |  |  |  |  |  | RNA2: | Complete |
|  | 6 | Maize associated bymovirus | Zea mays | SRR10376504 | 2175 | RNA1: | Complete |
|  |  |  |  |  |  | RNA2: | Complete |
|  | 7 | Zoysia associated bymovirus | Zoysia japonica | SRR9970175 | 5918 | RNA1: | Complete |
|  |  |  |  |  |  | RNA2: | Complete |
| **Ipomovirus** | 8 | Silene associated ipomovirus | Silene nutans | ERR4643771 | 1594 |  | Complete |
| **Macrophovirus** | 9 | Lophiostoma associated macrophovirus | Lophiostoma macrostomum | SRR1797204 | 795 |  | Complete |
|  | 10 | Puccinia associated macrophovirus | Puccinia trititica | SRR11929924 | 1321 |  | Complete |
|  |  |  | Puccinia striiformis | SRR621436 | 1272 |  | Complete |
|  |  |  | Triticum aestivum | SRR11563887 | 27262 |  | Complete |
|  | 11 | Maize associated macrophovirus | Zea mays | SRR12300538 | 943 |  | Complete |
|  | 12 | Lophodermella associated macrophovirus | Lophodermella | SRR15498770 | 9606 |  | Complete |
|  | 13 | Switchgrass associated macrophovirus | Zea mays | SRR15538946 | 2499 |  | Complete |
|  |  |  | Panicum virgatum | SRR10849187 | 73107 |  | Complete |
|  | 14 | Bat associated macrophovirus | Virome from bats | SRR2063913 | 2041 |  | Complete |
|  |  |  | Cheese microbiota | ERR6897493 | 1859 |  | Complete |
|  | 15 | Ovis associated macrophovirus | Feces from Ovis aries | ERR3143227 | 6228 |  | Near-complete |
|  | 16 | Anacyclus associated macrophovirus | Anacyclus depressus | SRR5237233 | 501 |  | Complete |
|  | 17 | Helianthus associated macrophovirus | Helianthus annuus | SRR12450951 | 645 |  | Complete |
|  | 18 | Cheese associated macrophovirus | Cheese microbiota | ERR6897690 | 609 |  | Near-complete |
| **Poacevirus** | 19 | Sugarcane associated poacevirus | Sugarcane and wild Poaceae | SRR14205968 | 56227 |  | Complete |
|  | 20 | Lolium associated poacevirus | Lolium multiflorum | SRR3100255 | 132426 |  | Complete |
|  |  |  | Festulolium braunii | SRR13346904 | 111093 |  | Complete |
| **Potyvirus** | 21 | Gentiana associated potyvirus | Gentiana atuntsiensis | SRR9835550 | 236824 |  | Complete |
|  |  |  | Gentiana microdonta | SRR9856849 | 95293 |  | Complete |
|  | 22 | Phragmites associated potyvirus | Phragmites karka | SRR9670022 | 386219 |  | Complete |
|  | 23 | Nardostachys associated potyvirus | Nardostachys jatamansi | SRR20715441 | 130555 |  | Complete |
|  | 24 | Senecio associated potyvirus | Senecio squalidus | SRR9326616 | 54523 |  | Complete |
|  | 25 | Angelica associated potyvirus | Angelica dahurica | SRR12919305 | 501767 |  | Complete |
|  | 26 | Scrophularia associated potyvirus | Scrophularia nodosa | SRR11906425 | 398367 |  | Complete |
|  | 27 | Gynostemma associated potyvirus | Gynostemma pubescens | SRR11573084 | 56697 |  | Complete |
|  | 28 | Taraxacum associated potyvirus | Taraxacum antungense | SRR10321514 | 387718 |  | Complete |
|  | 29 | Calante associated potyvirus | Calanthe hoshii | DRR172086 | 468567 |  | Complete |
|  | 30 | Misopates associated potyvirus | Misopates orontium | ERR10378007 | 50021 |  | Complete |
|  | 31 | Primula associated potyvirus | Primula veris | ERR5762859 | 449377 |  | Complete |
|  | 32 | Thymus associated potyvirus | Thymus quinquecostatus | SRR21700033 | 13098 |  | Complete |
|  | 33 | Silvianthus associated potyvirus | Silvianthus bracteatus | SRR12009658 | 350669 |  | Complete |
|  | 34 | Lancea associated potyvirus | Lancea tibetica | SRR11994215 | 12570 |  | Complete |
|  | 35 | Asparagus associated potyvirus | Asparagus officinalis | SRR1642915 | 2725 |  | Complete |
|  | 36 | Bupleurum associated potyvirus 1 | Bupleurum chinense | SRR17234910 | 79352 |  | Complete |
|  | 37 | Bupleurum associated potyvirus 2 | Bupleurum chinense | SRR19897009 | 28941 |  | Complete |
|  | 38 | Ranunculus associated potyvirus | Ranunculus auricomus | SRR21162053 | 27330 |  | Complete |
|  | 39 | Solanum associated potyvirus | Solanum lycopersicum | SRR21377516 | 2002 |  | Complete |
|  | 40 | Sedum associated potyvirus | Sedum alfredii | SRR7594980 | 63791 |  | Complete |

### Potential new members for the distinctive *Bymovirus* genus

In this study, we prioritized the characterization of viruses outside the extensively represented genus *Potyvirus* and its sister genus *Rymovirus*, aiming to uncover novel evolutionary trajectories and refine the taxonomic framework of the *Potyviridae* family. According to ICTV, the genus *Bymovirus* currently comprises six recognized species. Remarkably, our analyses enabled the identification and complete sequencing of the bipartite genomes of potentially four additional bymoviruses (Table 1), thereby substantially expanding the known diversity of this genus. The classification of these viruses within the genus *Bymovirus* is supported by both their phylogenetic placement based on NIb amino acid sequence comparisons (Figure 1) and their characteristic bipartite genome architecture (Figure 2A).

**Figure 1.**
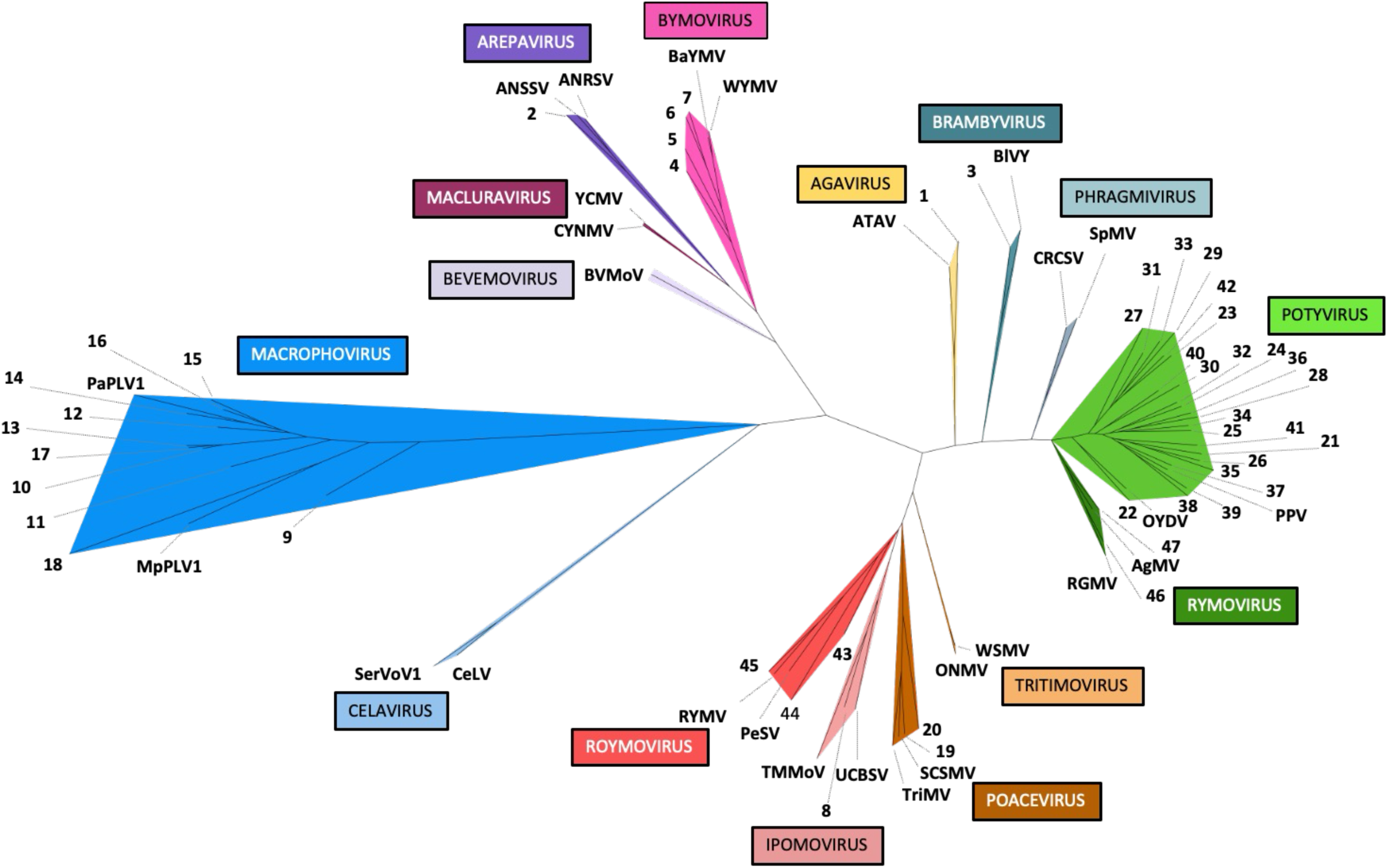
Phylogenetic tree of NIb replicase proteins from potyvirids. Genera are indicated by different colours. Newly identified viral isolates are labelled with numbers as shown in Table 1. When available, two previously reported species from each genus (see Supplementary Table 1 for details) were included to better define the placement of the different genera within the tree.

**Figure 2.**
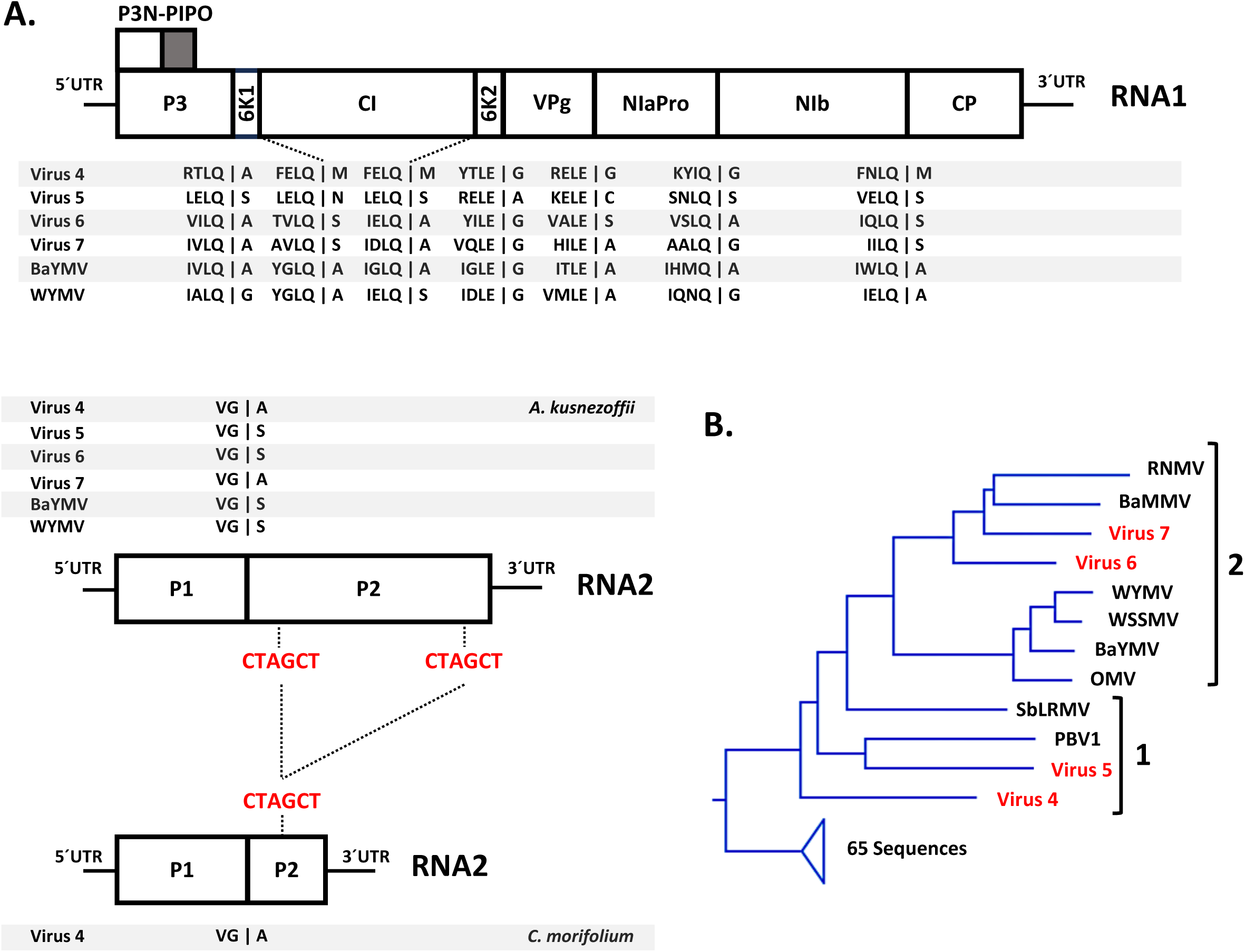
Analysis of bymovirus species. (A) Genomic organization of bymovirus isolates (Table 1, viruses 4-7) and the previously reported BaYMV and WYMV isolates (see Supplementary Table 1 for details), showing the two RNA segments and the predicted polyprotein cleavage sites for each species. Vertical bars indicate cleavage sites. Cleavage motifs for diverse viral proteases are shown below the schematic representation of viral genomes. In RNA2, a recombination event identified in virus 4 infecting *C. morifolium* is indicated, with repeated nucleotides highlighted in red. (B) Phylogenetic tree of NIb proteins from bymoviruses. The sequences included in the tree shown in Figure 1 were analysed together with previously reported bymovirus species (see Supplementary Table 1 for details). The two subclusters infecting Poaceae and non-Poaceae hosts are labelled 1 and 2, respectively.

As indicated in Table 1, virus 4 was detected in two different plants (*Aconitum kusnezoffii* and *Chrysanthemum morifolium*), displaying 98.6% and 96.7% nucleotide identity when comparing the coding sequences from RNA1 and RNA2 of the two variants, respectively. Notably, this comparison also revealed that the virus present in *C. morifolium* harbours a large deletion within the P2 coding sequence of RNA2 (Figure 2A, RNA2 panel). This feature is reminiscent of previously reported deletions in the RNA2 of other bymoviruses (Jacobi et al., 1995, Dessens et al., 1995, Kuhne et al., 2003, Zheng et al., 2002). Moreover, a detailed analysis of nucleotide sequences from both P2-intact and P2-deleted variants of virus 4 showed that the missing fragment is flanked by an identical six-nucleotide repeat (Figure 2A, RNA2 panel), supporting the hypothesis that the deletion arose via recombination during virus replication. Such deletions in the P2 coding sequence have been associated with loss of transmissibility by its vector, as they emerged after multiple passages by mechanical inoculation (Jacobi et al., 1995, Dessens et al., 1995, Zheng et al., 2002). However, this deletion has also been correlated with within-host fitness costs (You and Shirako, 2010). Therefore, we cannot rule out the possibility that these genome deletions in bymoviruses may represent a self-attenuation strategy that promotes long-term plant-virus coexistence, as recently reported for a distinct natural deletion in other potyvirids (Qin et al., 2025).

According to the ICTV, all officially recognized members of this genus infect monocotyledonous plants belonging to the Poaceae family, including barley, wheat, oat, and rice. As shown in Table 1, among the putative bymoviruses described here, only two were detected in Poaceae hosts (*Zea mays* and *Zoysia japonica*), whereas the remaining two were identified in samples derived from dicotyledonous plants (*A. kusnezoffii*, *C. morifolium*, and *Eutrema japonicum*). Although contamination with RNA from infected Poaceae plants cannot be entirely ruled out, infection of non-Poaceae hosts by bymoviruses would not be unexpected, as non-Poaceae plants have recently been reported to harbour newly discovered bymoviruses (Ohki et al., 2021, Cao et al., 2024). Notably, putative bymoviruses associated with non-Poaceae hosts form a basal subcluster in a NIb-based phylogenetic tree (subcluster 1), whereas the bymovirus infecting Poaceae plants groups within a distinct subcluster (subcluster 2) (Figure 2B). Together, these observations suggest that host range specialization in bymoviruses—particularly their adaptation to Poaceae hosts—may be linked to distinct evolutionary lineages.

### A potential second representative for the *Brambyvirus* genus

The genus *Brambyvirus* currently comprises only a single species according to ICTV, *Brambyvirus rubi* (blackberry virus Y, BlVY), which infects plants of the genus *Rubus* and encodes a type-B P1 [P1(b)] protein (Valli et al., 2011) containing an AlkB domain and a very short HCPro lacking typical silencing suppression-related domains as distinctive features. (Susaimuthu et al., 2008). Our phylogenetic analysis based on NIb assigned virus 3 (Table 1) to the genus *Brambyvirus* (Figure 1). This virus also shares distinctive features with BlVY, including the presence of a short HCPro and a zinc finger domain within its P1(b) (Figure 3A). Notably, the P1(b) protein, in contrast to type-A P1 [P1(a)], has been shown to act as RNA silencing suppressor in other viruses (Valli et al., 2006, Valli, 2007 #140, Giner et al., 2010, Tatineni et al., 2012, Young et al., 2012), suggesting that P1(b) may also fulfil this role in brambyviruses. Recognition sites for viral proteases appear to be conserved when comparing both viruses (Figure 3A). In this regard, it is worth noting that the cleavage sites FVQQ│N and SEPA│L, previously proposed by Susaimuthu and colleagues for processing BlVY polyprotein at the CI-6K2 and NIb–CP junctions, respectively, do not conform to the more commonly conserved (L/I)X(F/Y)(Q/E)│(A/S/G/N) motif for NIaPro-mediated processing (Figure 3A). In contrast, the alternative motif IDFQ│S, located in close proximity to the aforementioned sites, shows a much better match to the consensus. Notably, similar motifs are also present in virus 3 at the CI-6K2 and NIb-CP junctions (LTFQ│N and LTFQ│A, respectively) (Figure 3A). Based on these observations, we propose that IDFQ│S represents the actual NIaPro cleavage site responsible for processing at the CI-6K2 and NIb-CP junctions in BlVY (Figure 3A). Unfortunately, the assembly of the virus 3 genome does not include the 5’ end, which includes the N-terminal region of P1 and the 5’ UTR. This limitation precludes determining whether virus 3 also encodes an AlkB domain within P1 and, consequently, whether the presence of AlkB is a common feature of brambyviruses or a specific characteristic of BlVY (Figure 3A). Unlike BlVY, virus 3 was identified in *Forsythia suspensa* (Table 1), a host that is not related to *Rubus*.

**Figure 3.**
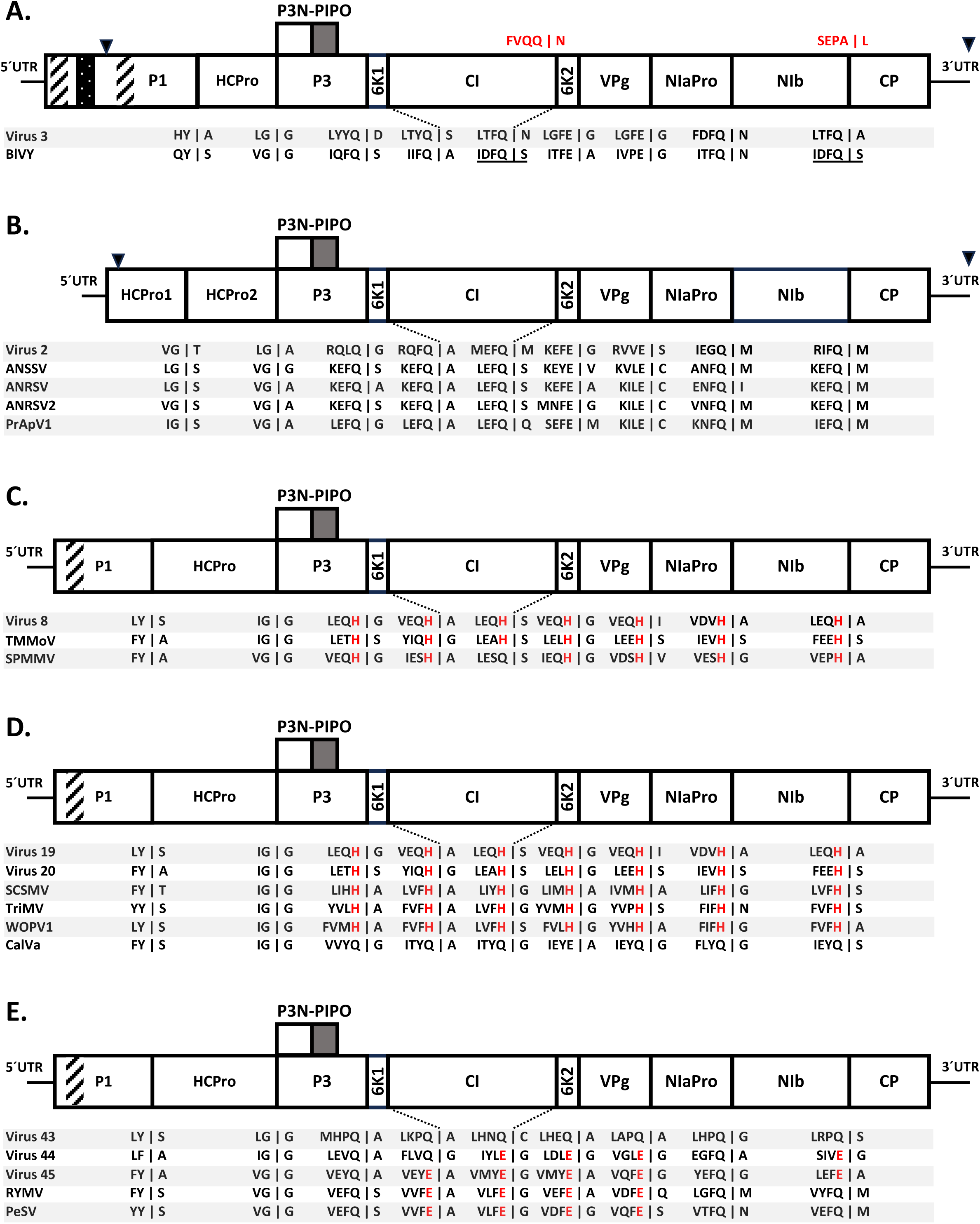
Genomic organization of selected potyvirid genera. (A-E) Genome organization of *Brambyvirus*, *Arepavirus*, *Ipomovirus*, *Tritimovirus* and *Roymovirus*, respectively. Vertical bars indicate predicted cleavage sites. Cleavage motifs for diverse viral proteases are shown below the schematic representation of viral genomes, and red letters highlight cleavage-related features that warrant special attention. Rectangles with diagonal bars indicate zinc-finger motifs, and a dotted rectangle mark the AlkB motif. Black triangles on top indicate the length of the reconstructed genomes in cases of incomplete assembly. Newly identified viruses are labelled with numbers as shown in Table 1, whereas previously reported members are included for cleavage-site comparison (see Supplementary Table 1 for details). Potentially incorrect previously reported cleavage sites in BlVY are shown in red.

### Another potential member of the *Arepavirus* genus

According to ICTV, the genus *Arepavirus* currently comprises only three viruses known to infect areca palm (order Arecales, family Arecaceae) in nature. Although the genome of virus 2 is not totally complete (Table 1), lacking the 5’ UTR and part of the HCPro1 coding sequence, phylogenetic analysis based on NIb proteins places this virus within the *Arepavirus* genus (Figure 1). In addition, it exhibits the characteristic genome organization observed in arepaviruses, namely the presence of two copies of HCPro in tandem (HCPro1-HCPro2) acting as leader proteases (Figure 3B). The protease recognition sites appear to be fairly conserved when comparing both viruses (Figure 3B). Remarkably, this virus was identified in *Vigna angularis* (order Fabales, family Fabaceae), a host unrelated to areca palm, suggesting a potential expansion of the host range within the genus. This notion is further supported by the recent identification of another member of this genus in *Psychotria rubra* (order Gentianales, family Rubiaceae), also a non-areca host (Thava Prakasa Pandian et al., 2024).

### New putative members of the *Ipomo*-, *Poace*-, *Roymo- and Tritimovirus* clade

As shown in Figure 1, our NIb-based phylogenetic analysis groups the *Ipomovirus*, *Poacevirus*, *Roymovirus*, and *Tritimovirus* genera within the same clade. This observation is consistent with phylogenetic trees based on CP (Supplementary Figure 1) and the complete polyprotein (see, for example, (Choi and Hahn, 2025)), suggesting a shared evolutionary origin. The genus *Ipomovirus* currently comprises seven recognized members according to ICTV, which display diverse genome organizations, particularly in the region encoding leader proteases. Specifically, two viruses encode a P1(b)-HCPro arrangement, three encode P1(a)-P1(b) proteases, whereas the remaining two encode only P1(b) (lacking HCPro) (Dombrovsky et al., 2014, Desbiez et al., 2016). Among the viruses identified in our in-silico survey, virus 8, found in a sample from *Silene nutans*, appears to represent a novel member of the *Ipomovirus* genus (Table 1). This is supported by the NIb-based phylogenetic analysis (Figure 1) and by its genome organization, which is consistent with the other members of this genus that encode a P1(b)-HCPro cassette (Figure 3C). Notably, this particular subgroup of ipomoviruses expressing P1(b)-HCPro, including virus 8, exhibits atypical NIaPro cleavage sites in which the conserved glutamine (Q) or glutamic acid (E) at the P1 position is replaced by histidine (H) (Figure 3C). This unusual substitution suggests a divergence in NIaPro substrate specificity, and may reflect an evolutionary adaptation of this ipomovirus subgroup, potentially associated with distinct regulatory requirements during polyprotein maturation.

The *Poacevirus* genus currently comprises four recognized members according to ICTV, all of which share a conserved genome organization characterized by the presence of P1(b)-HCPro as leader proteases (Figure 3D). Our in-silico survey identified two putative new members, virus 19 and virus 20, detected in samples from plant species belonging to the Poaceae family, such as *Lolium multiflorum*, *Festulolium braunii*, sugarcane and its wild relatives (Table 1). Some poaceviruses are characterized by unusually large 5’ untranslated regions (UTRs), including *Poacevirus tritici* (triticum mosaic virus, TriMV), *Poacevirus avenae* (wild oat poacevirus 1, WOPV1), and *Poacevirus caladeniae* (caladenia virus A, CalVA), whose 5’ UTRs are 739, 531, and 366 nt long, respectively (Tatineni et al., 2009, Huang et al., 2024, Wylie et al., 2012). The assembled genomes of virus 19 and virus 20 also contain exceptionally long 5’ UTRs exceeding 650 nt; however, their precise lengths would need to be experimentally determined by 5’ RACE. Importantly, the genomic regions of virus 19 and virus 20 that show the highest nucleotide-level similarity to other poaceviruses are particularly located within a large segment of their respective 5’ UTRs (Supplementary Figure 2). In the case of virus 19, a region spanning 441 nt within the 5’ UTR exhibits 74% nucleotide identity to the corresponding region of TriMV (Supplementary Figure 2A). Similarly, virus 20 contains a 393-nt segment in its 5’ UTR that aligns with 75 % nucleotide identity to the corresponding region of Poaceae Liege poacevirus (GenBank accession number ON137719), a putative member of the genus not yet officially recognized by the ICTV, whose 5’ UTR has 689 nt (Supplementary Figure 2B). Together with previously published data on the role of TriMV 5’ UTR on promoting and enhancing cap-independent translation (Roberts et al., 2015, Jaramillo-Mesa et al., 2019), this unexpected level of conservation across extensive portions of the 5’ UTR among different viruses further supports the notion that this region plays critical roles in viral infection. Finally, it is worth noting that, with the exception of CalVA—the only member identified in a non-poaceous host (Wylie et al., 2012)—all other poaceviruses examined also exhibit histidine (H) at the P1 position of NIaPro cleavage sites (Figure 3D), a feature likewise observed in a distinct subgroup of ipomoviruses (see above).

With only two species currently recognized by the ICTV, *Roymovirus* is the most underrepresented genus within this *Potyviridae* clade. Our pipeline identified three particular viruses, virus 43, virus 44, and virus 45, which appear to belong to the genus *Roymovirus*, as indicated by the NIb-based phylogenetic analysis (Figure 1). In addition, these viruses display the same genome organization as the species previously assigned to this genus by the ICTV (Figure 3E). Moreover, with the exception of virus 43, which contains glutamine (Q), glutamic acid (E) is overrepresented at the P1 position of the NIaPro cleavage sites (Figure 3E). Although virus 44 appears to share part of its host range with *Roymovirus rosae* (rose yellow mosaic virus, RYMV), as both infect hosts from the Rosaceae family (ICTV, and Table 1), an analysis of the plant species from which both officially recognized (ICTV) and newly identified roymoviruses were isolated (Table 1) indicates that the overall host range of roymoviruses is remarkably broad.

### *Agavirus*, a potential new genus with two members within the family *Potyviridae*

Our pipeline identified a novel member of the family *Potyviridae*, hereafter referred to as virus 1, in a sample from *Acrocomia aculeata* (order Arecales, family Arecaceae) (Table 1). BLASTP analysis using the full amino acid sequence of virus 1 yielded only potyvirid hits with very low sequence identity. The most similar virus (35% amino acid identity) was agave tequilana agavirus (ATAV), which has not yet been officially recognized by the ICTV. To our knowledge, no peer-reviewed study has been published on ATAV; however, its genome sequence has been deposited in the NCBI database (GenBank accession number MZ682615) with the indication that it may represent a novel genus within the family *Potyviridae*. Consistent with this annotation, NIb- and CP-based phylogenetic analyses showed that virus 1 and ATAV form a well-defined clade that is clearly independent of all currently recognized genera within the family *Potyviridae* (Figure 1 and Supplementary Figure 1). As observed for most members of the family, NIaPro-mediated proteolytic processing of viral polyproteins depends on the presence of glutamine (Q) or glutamic acid (E) at the P1 position (Figure 4A). Notably, the majority of these cleavage sites also share alanine at the P4 position (Figure 4A). We propose that these viruses comprise a new genus within the family *Potyviridae*. In recognition of the researchers who determined the full-length genome sequence of ATAV and originally proposed a potential name for this putative new genus, we also propose the name *Agavirus* for this new group.

**Figure 4.**
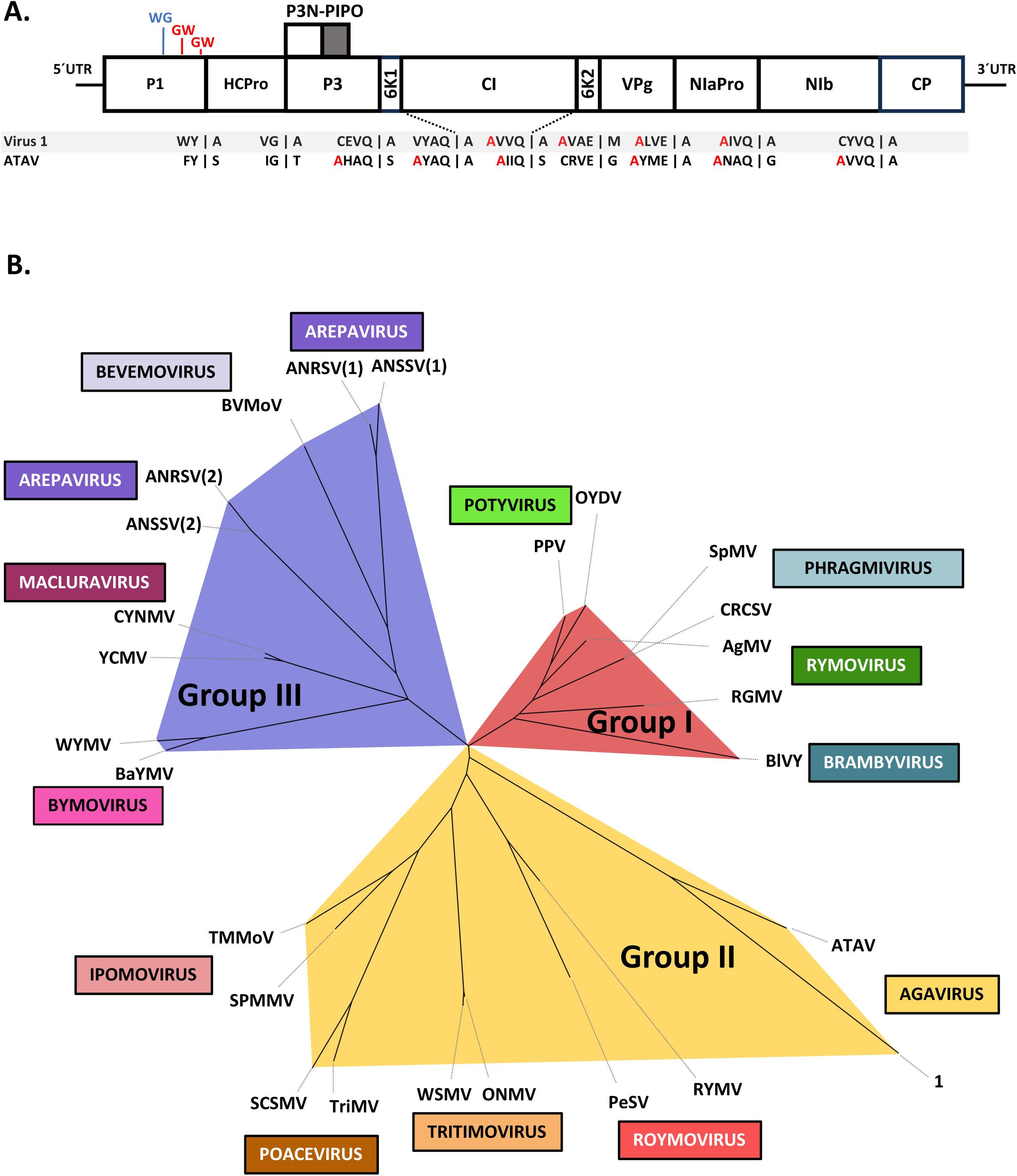
Analysis of the proposed genus *Agavirus*. (A) Genome organization of virus 1 identified in this study (Table 1) and the previously reported ATAV (see Supplementary Table 1 for details). Vertical bars indicate predicted cleavage sites. Cleavage motifs for diverse viral proteases are shown below the schematic representation of viral genomes, and red letters highlight the unusual amino acid residue at the P4 position. GW/WG motifs shown in red are present in both agaviruses, whereas the motif shown in blue is only present in ATAV. (B) Phylogenetic tree based on the HCPro protease domains of *Potyviridae* species (see Supplementary Table 1 for details). The three groups are shown in different colours.

Evidence supporting the placement of these viruses in a novel genus is not limited to phylogenetic inference. For instance, the HCPro proteins encoded by agaviruses are remarkably small (264 and 265 amino acids for virus 1 and ATAV, respectively) and lack the conserved FRNK motif, which is known to play a central role in RNA silencing suppression in potyviruses (Valli et al., 2018). Although these two features are characteristic of a specific group of HCPro proteins expressed by arepa-, bymo-, bevemo-, and macluraviruses, collectively referred to as group III HCPro factors (Qin, 2020), phylogenetic analysis of the conserved cysteine protease domain of potyvirid HCPro proteins instead supports the classification of agavirus HCPro factors within group II (Figure 4B). This group comprises HCPro proteins encoded by members of the ipomo-, poace-, roymo-, and tritimovirus clade, and the placement of agaviral HCPro proteins within this group is consistent with (i) the close phylogenetic proximity of the agavirus clade to ipomo-, poace-, roymo-, and tritimoviruses in the NIb- and CP-based phylogenetic trees (Figure 1 and Supplementary Figure 1), and (ii) the fact that agaviruses encode a P1 leader protease (Figure 4A), as do members of these genera. Remarkably, group II HCPro factors are characterized by a lack of RNA silencing suppression activity, and viruses encoding these HCPros instead rely on P1 to counteract RNA silencing. Therefore, it is intriguing to find that agavirus P1 proteins do not contain the characteristic zinc-finger motif, which has been linked to RNA silencing suppression activity in ipomo-, poace-, roymo-, and tritimoviruses (Valli et al., 2008). Together, the absence of conserved motifs implicated in RNA silencing suppression in both P1 and HCPro raises the question of which viral protein mediates RNA silencing suppression in agaviruses. It remains possible that agavirus HCPro proteins counteract RNA silencing-based antiviral defenses through non-canonical or as-yet-unknown mechanisms, as has been reported for small HCPros belonging to group III (Qin et al., 2020, Hu et al., 2023, Chen et al., 2023). Alternatively, agavirus P1 proteins may participate in interactions with components of the RNA silencing machinery, as they harbour GW/WG motifs (Figure 4A), which have been generally associated with ARGONAUTE binding, as in the case of *Ipomovirus lenisbatatae* (sweet potato mild mottle virus) (Giner et al., 2010). Experimental validation will be required to determine the contribution of these agavirus proteins to RNA silencing suppression.

### A novel group of potyvirids is associated with fungi and oomycetes

Our pipeline enabled the assembly of complete or near-complete genomes of ten previously undescribed viruses belonging to the family *Potyviridae* (hereafter referred to as viruses 9–18), detected across multiple samples from diverse origin (Table 1). BLASTP analysis using the full-length polyprotein sequences revealed that these viruses do not cluster with any of the potyvirid genera currently recognized by the ICTV, sharing only low levels of amino-acid identity with established members of the family. Instead, the closest matches corresponded to a small number of highly divergent potyvirid-like viruses, including three viruses previously reported in the literature and associated with fungal and oomycete samples—macrophomina phaseolina poty-like virus 1 (MpPLV1) (Wang et al., 2021), plasmopara-associated poty-like virus 1 (PaPLV1) (Chiapello et al., 2020), and uromyces potyvirus A (UrPA) (Jo et al., 2022)—as well as some unclassified poty-related sequences deposited in the NCBI database, none of which has so far been assigned to an official genus. Consistent with these BLASTP-based observations, NIb-based phylogenetic analyses showed that viruses 9-18, together with MpPLV1 and PaPLV1, cluster in a well-supported and deeply divergent clade that is clearly separated from all currently recognized genera within the family *Potyviridae* (Figure 1). Moreover, both NIb- and CP-based phylogenetic analyses indicate that this lineage is most closely related to the genus *Celavirus* (Figure 1 and Supplementary Figure 1), which has been considered the most divergent group of potyvirids to date (Choi and Hahn, 2025). Collectively, these results strongly support the existence of a highly divergent and previously unrecognized lineage of potyvirids. In light of the initial hypothesis proposed by Wang and collaborators that potyvirids may infect fungal hosts, following the identification of a poty-like virus in the fungus *Macrophomina phaseolina* (Wang et al., 2021), we propose the name *Macrophovirus* for this novel group.

It is worth noting that three of the ten macrophoviruses listed in Table 1 have been identified previously. To our knowledge, the sequences corresponding to viruses 10 and 13 have not been described in peer-reviewed publications; however, highly similar sequences are available in the NCBI database as *Puccinia triticina*–associated virus 4 (GenBank accession number PX650069) and Switchgrass poty-like virus 1 (GenBank accession number PP996020), respectively. Virus 17 shares 99% amino-acid identity with the previously reported UrPA (GenBank accession number MK231047) (Jo et al., 2022). Notably, unlike the original UrPA sequence, the genome of virus 17 is complete and includes the 5′ untranslated region and the 5′ end of the first cistron, which were absent from the reported UrPA sequence.

Based on the collected sequence data, macrophoviruses possess relatively small genomes, ranging from approximately 6.9 kb, as observed for virus 9, to 8.1 kb, as in the case of virus 12. Basic CD-Search analyses, which typically detect conserved potyvirid domains in most potyviruses (Supplementary Figure 3A), revealed that macrophoviruses encode a CI-like helicase, an NIaPro-like C4 protease, and an NIb-like RNA-dependent RNA polymerase (RdRP), arranged at genomic positions similar to those observed in other members of the family *Potyviridae* (Supplementary Figure 3B). This information enabled us to infer approximate cistron boundaries and, consequently, to predict potential NIaPro cleavage sites. Notably, these sites conform to the loosely conserved motif (R/K)-X-(F/Y/W)-(Q/E)│(A/G/S/N/M). Based on this information, we identified several putative mature factors generated during infection through NIaPro-mediated processing (Figure 5A). Specifically: (i) the terminal cistron encodes the coat protein (CP), as is typical for potyvirids, and it shows limited similarity to the CP of celaviruses (Supplementary Figure 1); (ii) the cistron encoding the CI-like helicase is shorter than that of canonical potyvirids because it lacks the coding sequence for the conserved Poty-PP domain (Supplementary Figure 3); (iii) a P3-like cistron is present, which would encode a factor lacking detectable primary sequence similarity with known potyvirid P3 proteins but appears to encode a transmembrane domain, as observed for standard potyvirid P3 factors; notably, (iv) this P3 cistron does not contain the conserved overlapping ORF encoding PIPO; (v) small peptide-coding sequences are predicted to flank the CI cistron, although they show no detectable similarity to 6K1 or 6K2; and (vi) additional small peptide-coding cistrons may also be present (Figure 5A).

**Figure 5.**
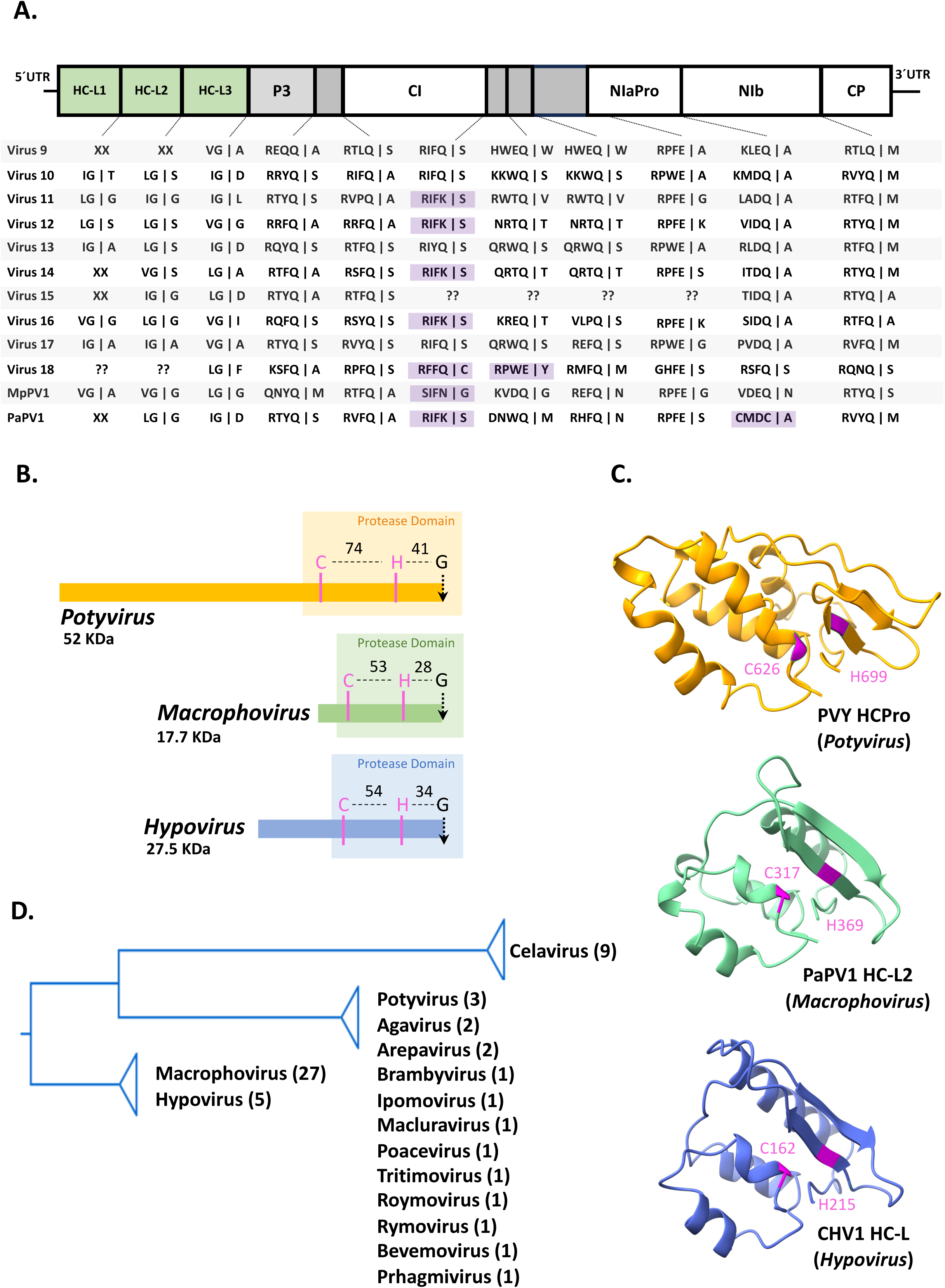
Analysis of the proposed genus *Macrophovirus*. (A) Genome organization of the newly identified viral species (viruses 9-18; Table 1) and the previously reported MpPV1 and PaPV1 (see Supplementary Table 1 for details). Vertical bars indicate predicted cleavage sites. Cleavage motifs for diverse viral proteases are shown below the schematic representation of viral genomes. Motifs highlighted in purple indicate unusual or absent cleavage sites. “XX” indicates absence of the corresponding cistron, whereas question marks indicate genome segments that were not assembled from the RNAseq data. Only proteins showing clear homology to potyvirid proteins are labelled on the genome. HCPro-like (HC-L) proteins are shown as green boxes. (B) Schematic diagrams of the HCPro from *Potyvirus* (PVY), *Macrophovirus* (PaPV1) and *Hypovirus* (CHV1) (see Supplementary Table 1 for details). The catalytic residues C and H are shown in pink, and the cleavage residue G is shown in black; the distances between residues are indicated in amino acids. (C) Three-dimensional models of protease motifs from proteins shown in (B). Catalytic residues (C and H) are highlighted in pink. (D) Phylogenetic tree including the HCPro and HC-L protease domains from potyvirids and the hypoviruses CHV1, HVC and MKrcHV2 (see Supplementary Table 1 for details).

A careful inspection of the 5′ genomic region of macrophoviruses revealed the presence of leader proteases, as observed in most members of the family *Potyviridae*. Specifically, we identified cysteine proteases related to HCPro that are arranged in variable configurations, ranging from one to three tandem copies. Although this represents the first report of a three-cysteine-protease tandem arrangement in potyvirids, the most striking observation is that these proteases display greater similarity to leader papain-like proteases from the family *Hypoviridae* (Koonin et al., 1991), a group of +ssRNA viruses infecting fungi (Chiba et al., 2023). Remarkably, although not reported in the literature, we found some examples of hypoviruses also exhibiting tandem arrangements of multiple cysteine proteases at the N-terminus of the viral polyprotein (Supplementary Figure 4).

Consistent with that similarity with hypoviruses, these HCPro-like proteases (hereafter referred to as HC-L) of macrophoviruses are substantially smaller than canonical HCPro proteins found in other potyvirids. Moreover, the spacing between the catalytic cysteine (C) and histidine (H) residues, as well as the distance to the conserved glycine (G) residue at the cleavage site, is markedly reduced (Figure 5B). This compact organization closely mirrors that observed in HC-L from hypoviruses and contrasts with the larger inter-residue distances characteristic of canonical potyvirid HCPro proteins. Predicted three-dimensional structures indicate that macrophoviral and hypoviral HC-L factors adopt highly similar folds (Figure 5C). Consistently, a structural similarity search using the model of PaPV1 HC-L2 as template identified CHV1 HC-L as a close structural match. Phylogenetic analysis incorporating canonical HCPro proteins from potyvirids and HC-L proteins from macrophoviruses and hypoviruses further supports this distinction, with all HC-L factors clustering into a separate, well-supported branch that is clearly distinct from the clade encompassing canonical potyvirid HCPro proteins (Figure 5D). As reported for P29 (an HC-L protein) from *Alphahypovirus cryphonectriae* (cryphonectria hypovirus 1, CHV1) (Suzuki et al., 2003, Segers et al., 2006), it is expected that HC-L factors from macrophoviruses counteract host RNA-silencing-based antiviral activities.

Finally, we identified an unusual genomic organization in virus 18, as the sequence encoding the CP is not part of the main polyprotein reading frame. Although the CP cistron is located at the canonical position near the 3′ end of the potyvirid genome, it is encoded in an alternative reading frame relative to the primary ORF (Supplementary Figure 3C). Importantly, this genomic arrangement was independently observed in contigs assembled from two distinct samples (ERR6897690 and ERR6897493, Table 1), strongly arguing against assembly artifacts and supporting its biological relevance. Remarkably, the main reading frame retains sufficient coding capacity in this region to potentially encode an additional protein, suggesting that this genomic architecture may allow the expression of both the CP and an extra factor through distinct translational strategies (Supplementary Figure 5). CP expression in this context is therefore likely achieved via a noncanonical mechanism, such as leaky ribosomal scanning, programmed ribosomal frameshifting, polymerase slippage, internal translation initiation, or the production of a specific subgenomic RNA. This arrangement may reflect selective pressure to uncouple CP expression from polyprotein processing while simultaneously accommodating an accessory protein in the main ORF, thereby increasing the functional versatility and regulatory complexity of the viral genome.

Taken together, the detection of several of these viruses in pure fungal samples, fungus- and oomycete-infected plants, fungus-containing food products, and/or soil ((Wang et al., 2021, Chiapello et al., 2020, Jo et al., 2022); and Table 1), along with the presence of hypovirus-like leader proteases, the absence of plant cell-to-cell movement factors (Figure 5), and the emergence of genomic and proteomic features not previously described in potyvirids, strongly supports the hypothesis that macrophoviruses infect fungi and oomycetes in nature and represent the most divergent lineage within the family *Potyviridae*.

## DISCUSSION

The large-scale mining of publicly available RNA-seq data conducted in this study substantially expands the known diversity of the family *Potyviridae* and reveals unexpected evolutionary trajectories within this group. Beyond the identification of multiple novel species across established genera, our analyses uncovered a deeply divergent lineage of potyvirids associated with fungal and oomycete samples, for which we propose the genus *Macrophovirus*. These viruses exhibit a combination of genomic, proteomic, and phylogenetic features that distinguish them from canonical plant-infecting potyvirids and strongly suggest adaptation to fungal and oomycete hosts. Together with the discovery of unusual leader proteases and atypical genomic organizations, our findings provide new insights into potyvirid evolution, host range expansion, and functional plasticity.

### HCPro and its relatives

Leader proteases play central roles in potyvirid infections, coordinating polyprotein processing and modulating host antiviral responses, particularly RNA silencing. In most potyvirids, these functions are carried out by HCPro, a multifunctional cysteine protease that acts as a potent suppressor of RNA silencing in plants (Valli et al., 2018). The discovery of HCPro-related leader proteases in macrophoviruses was therefore anticipated; however, their architecture and evolutionary origin proved highly unexpected. Macrophoviral leader proteases are arranged in variable tandem configurations, ranging from one to three copies, representing the first report of a triple cysteine-protease arrangement in potyvirids. More strikingly, these HC-L show limited similarity to canonical potyvirid HCPro factors but instead resemble leader papain-like proteases of hypoviruses. The functional implications of this resemblance are substantial. In hypoviruses, leader papain-like proteases, such as P29 from CHV, are well-established suppressors of RNA silencing in their fungal hosts and even retain activity in heterologous plant systems (Suzuki et al., 2003, Segers et al., 2006). By analogy, macrophoviral HC-L factors are strong candidates to counteract RNA-silencing-based antiviral defenses in fungi and oomycetes. The presence of tandem HC-L copies may reflect duplication events that enabled functional specialization or increased robustness of silencing suppression. Interestingly, although celaviruses were previously thought to lack recognizable leader proteases (Choi and Hahn, 2025, Rose et al., 2019), our comprehensive reanalysis revealed the presence of HCPro-like proteins in this group, which are also arranged in tandem in the genomes of several celaviruses (Supplementary Figure 6). Phylogenetically, these celaviral HCPro-like proteins cluster together with canonical potyvirid HCPro factors (Figure 5D), yet they form a distinct, well-supported lineage, suggesting that they constitute an additional evolutionary group within the diversity of potyvirid HCPro proteins (Figure 4B).

Together, these observations indicate that leader proteases in potyvirids have undergone extensive diversification, likely driven by host-specific selective pressures and repeated functional reinvention.

### NIaPro cleavage sites

Proteolytic processing by NIaPro is essential for the maturation of potyvirid polyproteins; however, as an increasing number of divergent members of the family are being discovered, it has become evident that the sequence determinants governing NIaPro substrate specificity are remarkably flexible rather than strictly conserved. Our comparative analyses reveal substantial variability in NIaPro cleavage motifs across potyvirids, particularly at the P1 position, where substitutions of glutamine—the most common amino acid at this position (Adams et al., 2005a)—are frequently observed. This plasticity is especially pronounced in specific lineages, notably within the cluster comprising ipomo-, poace-, roymo-, and tritimoviruses.

Accurate identification of cleavage sites is obviously critical for defining mature viral proteins and interpreting their functions. The extensive alignments presented here allowed us to refine cleavage site predictions and, in some cases, to reassess previously proposed processing boundaries. Importantly, imperfect or inefficient cleavage could give rise to partially processed fusion proteins, as previously observed (e.g., (Valli et al., 2022)), which may possess functions distinct from those of their fully cleaved counterparts (Revers and García, 2015). Such fusion proteins could expand the functional repertoire of the virus without increasing genome size, a strategy likely favored in compact RNA genomes. Experimental validation of cleavage efficiency and processing intermediates will be essential to fully understand how proteolytic regulation shapes infection cycles in potyvirids.

### 6K1, 6K2 and all the small things

The small membrane-associated proteins traditionally referred to as 6K1 and 6K2 were historically defined based on their approximate molecular weight and conserved position within the potyvirid polyprotein. However, as the diversity of the family *Potyviridae* has expanded beyond the *Potyvirus* genus, it has become increasingly clear that these proteins exhibit considerable size variability across different genera, with reported lengths ranging from approximately 7 to 14 kDa, even among members of a single genus such as *Bymovirus* (Wagh et al., 2016). Importantly, this variability in size does not necessarily reflect fundamental differences in structure or function. We argue that nomenclature based strictly on molecular weight (e.g., renaming these factors as “7K” or “14K”) is neither informative nor conceptually robust. Instead, proteins occupying equivalent positions in the polyprotein and sharing structural features—most notably the presence of transmembrane domains and association with replication membranes—should continue to be regarded as functional homologues of 6K1 and 6K2, regardless of their precise length. This approach better reflects evolutionary continuity and avoids artificial inflation of protein categories driven just by size differences.

In macrophoviruses, short peptide-coding regions flank the CI cistron, but they show little or no similarity to canonical 6K1 or 6K2 proteins. Whether these peptides represent highly divergent analogues of 6K proteins or entirely novel functional elements remains an open question. Given the increasing recognition that small viral peptides can play key roles in replication, membrane remodeling, host interaction and virus evolution (Dolja and Koonin, 2018), macrophoviruses may exemplify an extreme case of proteomic minimization and further innovation.

Overall, these observations highlight the need to reconsider rigid size-based classifications of small potyvirid proteins and to prioritize positional, structural, and functional criteria instead. Doing so will be essential for meaningfully comparing divergent potyvirids and for understanding how small proteins contribute to viral adaptation across markedly different hosts, including fungi and oomycetes.

### The chicken or the egg in *Potyviridae* evolution

One of the most intriguing questions raised by our findings concerns the evolutionary origin of potyvirids. Did potyvirids arise as plant viruses that later expanded their host range to fungi, or do macrophoviruses represent remnants of an ancestral fungal-infecting lineage from which plant-infecting potyvirids later emerged? Several observations lend support to the latter hypothesis. Macrophoviruses occupy a deeply divergent phylogenetic position within the family and display simplified or altered versions of hallmark potyvirid features, including compact genomes, reduced CI domains, absence of PIPO, and HC-L proteases. These features may reflect ancestral states retained in fungal hosts, whereas plant-infecting potyvirids may have acquired additional domains and functions to cope with the challenges of multicellular plant hosts, such as cell-to-cell movement and complex immune responses. Conversely, it is also possible that macrophoviruses evolved from plant-infecting ancestors through reductive evolution following a host jump to fungi. Under this scenario, the loss of movement proteins and the convergence toward HC-L proteases would represent adaptations to the fungal intracellular environment. A third scenario is that an ancestral potyvirid-like virus infected early eukaryotes prior to the divergence of plants, fungi, and oomycetes, with subsequent lineage-specific specialization and recurrent host jumps shaping the present-day diversity of the family. Distinguishing among these scenarios will require broader sampling of fungal-associated potyvirids and comparative analyses integrating molecular clocks, host ecology, and experimental studies of virus-host interactions.

Regardless of their evolutionary origin, macrophoviruses demonstrate that the host range of potyvirids is broader than previously appreciated, and that fungal and oomycete infections are not an evolutionary dead end but a viable and potentially ancient lifestyle for members of this family.

## ACKNOWLEDGMENTS

AI-assisted language editing tools were used for linguistic refinement. We are thankful to Juan Antonio García and Juan José López-Moya (the “*Polyviridie”* group) for fruitful discussions. This work was supported by grants PID2022-139314OB-I00 (to AAV) and PID2022-138530OB-I00 (to CSM) funded by MICIU/AEI/10.13039/501100011033 and by FEDER.

## Supplementary Figure

**Supplementary Figure 1. Phylogenetic tree of CP from potyvirids.** Genera are indicated in different colours. When available, two previously reported species from each genus (see Supplementary Table 1 for details) were included. The newly identified virus 1 (Table 1) was also included to better define the position of agaviruses together with ATAV.

**Supplementary Figure 2. Sequence conservation at the 5’ UTR of novel members of the genus Poacevirus.** (A) BLASTN analysis (NCBI) of the genome from virus 19 (Table 1). Nucleotide similarity to *Poacevirus tritici* (triticum mosaic virus, TriMV) isolate 19HM1 within the 5’ UTR and the CI and NIb cistrons is indicated with coloured boxes showing the corresponding percentage identities. The sequence alignment corresponding to the 5’ UTR region of virus 19 and TriMV is also shown. (B) BLASTN analysis (NCBI) of the genome from virus 20 (Table 1). Nucleotide similarity to Poaceae Liege poacevirus (PoLiV) isolate Latinne within the 5’ UTR and the CI and NIb cistrons is indicated with coloured boxes showing the corresponding percentage identities. The sequence alignment corresponding to the 5’ UTR region of virus 20 and PoLiV is also shown. The position of the start codon (AUG) and approximate genomic coordinates are indicated.

**Supplementary Figure 3. Comparison of conserved domains detected in a newly identified potyvirus and a newly identified macrophovirus.** Schematic representation of the conserved domains identified using the CD-SEARCH tool (NCBI), including their corresponding descriptions and E-values, based on the polyprotein sequences of (A) virus 36 and (B) virus 10 (Table 1). Polyprotein coordinates are indicated, and the location of the CI region within the polyprotein is highlighted in red.

**Supplementary Figure 4. Genomic organization of selected hypoviruses.** Shared genomic features of HVc, MpHV1, PoFHV2 and BFAHV (see Supplementary Table 1 for details) are highlighted in colour. Vertical bars indicate predicted polyprotein cleavage sites. Cleavage motifs for diverse viral proteases are shown below the schematic representation of viral genomes, whereas “XX” indicates absence of the corresponding cistron.

**Supplementary Figure 5. Genomic organization of newly identified Virus 18.** Genome organization of virus 18 (*Macrophovirus*), with predicted cleavage sites indicated by vertical bars. Cleavage motifs for diverse viral proteases are shown below the schematic representation of viral genomes. Motifs highlighted in purple indicate unusual or absent cleavage sites. The curve line indicates that the genomic sequence is incomplete. Only proteins showing clear homology to known potyvirid proteins are labelled on the genome. The CP is encoded outside the main ORF, and a protein of unknown function is encoded downstream of the NIb cistron.

**Supplementary Figure 6. Genomic organization of leader proteins within the genus *Celavirus*.** The 5’ end segment of all previously reported celaviruses are represented (see Supplementary Table 1 for details). Predicted cleavage sites are indicated by vertical bars; when a cleavage is uncertain, a question mark is shown. Catalytic residues are highlighted in red (H, D and S for P1; C and H for HCPro). The IxFG motif is indicated at the N-terminus of P1. The zinc-finger motif is indicated with a green dotted square. A red dotted rectangle indicates a signal peptide, and a rectangle with vertical bars indicate a putative protease cofactor.

