## Supplemental Table 1 for "Discovery of novel members of the *Potyviridae* family reveals expanded diversity, a broad host range, and evidence of fungal and oomycete infections"

**Supplementary Table S1: viruses used for this study.**

| Genus | Species | Binomial Name | Acronym | NCBI Accession |
|---|---|---|---|---|
| **Agavirus** | Agave tequilana agavirus | | ATAV | UYD62333 |
| **Arepavirus** | Areca palm necrotic ringspot virus | *Arepavirus arecae* | ANRSV | QZC92263 |
| | Areca palm necrotic spindle-spot virus | *Arepavirus arecamaculatum* | ANSSV | YP_009553653 |
| | Areca palm necrotic ringspot virus 2 | *Arepavirus karnatakense* | ANRSV2 | PQ197196 |
| | Psychotria rubra arepavirus 1 | | PrApV1 | BK068283 |
| **Bevemovirus** | Bellflower veinal mottle virus | *Bevemovirus campanulae* | BVMoV | YP_009508455 |
| **Brambyvirus** | Blackberry virus Y | *Brambyvirus rubi* | BlVY | YP_851006 |
| **Bymovirus** | Barley yellow mosaic virus | *Bymovirus hordeiluteum* | BaYMV | NP_148999 (RNA1) |
| | Wheat yellow mosaic virus | *Bymovirus triticitessellati* | WYMV | NP_059449 (RNA1) |
| | Rice necrosis mosaic virus | *Bymovirus oryzae* | RNMV | YP_009175089 (RNA1) |
| | Barley yellow mosaic virus | *Bymovirus hordeiluteum* | BaMMV | NP_604491 (RNA1) |
| | Wheat spindle streak mosaic virus | *Bymovirus tritici* | WSSMV | QGT40987 (RNA1) |
| | Oat mosaic virus | *Bymovirus avenae* | OMV | NP_659025 (RNA1) |
| | Soybean leaf rugose mosaic virus | | SbLRMV | BBD13978 (RNA1) |
| | Pea bymovirus 1 | | PBV1 | C_AA058313 (RNA1)* |
| **Celavirus** | Celery latent virus | *Celavirus apii* | CelV | YP_010087166 |
| | Serapias vomeracea-associated virus 1 | | SerVoV1 | DBB27949 |
| | Chrysanthemum x morifolium virus 1 | | ChrMoV1 | DBB27942 |
| | Dalzellia ubonensis virus 1 | | DalUbV1 | DBB27943 |
| | Leucadendron cela-like virus 1 | | LeuClV1 | DBB27944 |
| | Striga potyvirus B | | SalPlV2 | QVG60634 |
| **Ipomovirus** | Uganda cassava brown streak virus | *Ipomovirus manihotis* | UCBSV | YP_004063982 |
| | Tomato mild mottle virus | *Ipomovirus lycopersici* | TMMoV | YP_009509087 |
| | Sweet potato mild mottle virus | *Ipomovirus lenisbatatae* | SPMMV | OM471971 |
| **Macluravirus** | Chinese yam necrotic mosaic virus | *Macluravirus dioscoreachinense* | CYNMV | ANJ16151 |
| | Yam chlorotic mosaic virus | *Macluravirus dioscoreaflavitessellati* | YCMV | ASJ27536 |
| **Macrophovirus** | Macrophomina phaseolina poty-like virus | | MpPVL1 | QOE55591 |
| | Plasmopara viticola lesion associated poty-like virus 1 | | PaPLV1 | QHD64737 |
| **Phragmivirus** | Common reed chlorotic stripe virus | *Phragmivirus phragmii* | CRCSV | YP_009408143 |
| | Spartina mottle virus | *Phragmivirus spartinae* | SpMV | QVX19873 |
| **Poacevirus** | Sugarcane streak mosaic virus | *Poacevirus sacchari* | SCSMV | YP_006423941 |
| | Triticum mosaic virus | *Poacevirus tritici* | TriMV | XLG22950 |
| | Caladenia virus A | *Poacevirus caladeniae* | CalVA | YP_006666511 |
| | Wild oat poacevirus 1 | *Poacevirus avenae* | WOPV1 | PQ561517 |
| **Potyvirus** | Onion yellow dwarf virus | *Potyvirus cepae* | OYDV | NP_871002 |
| | Plum pox virus | *Potyvirus plumpoxi* | PPV | NP_040807 |
| **Roymovirus** | Passiflora edulis symptomless virus | *Roymovirus passifloralatentis* | PeSV | YP_010088106 |
| | Rose yellow mosaic virus | *Roymovirus rosae* | RYMV | YP_006908987 |
| **Rymovirus** | Agropyron mosaic virus | *Rymovirus agropyronis* | AgMV | YP_054400 |
| | Ryegrass mosaic virus | *Rymovirus lolii* | RGMV | NP_044727 |
| **Tritimovirus** | Oat necrotic mottle virus | *Tritimovirus avenae* | ONMV | NP_932608 |
| | Wheat streak mosaic virus | *Tritimovirus tritici* | WSMV | NP_046741 |
| | Tall oatgrass mosaic virus | *Tritimovirus arrhenatheri* | TOgMV | KF260962 |
| | Brome streak mosaic virus | *Tritimovirus bromi* | BrSMV | Z48506 |
| | Wheat eqlid mosaic virus | *Tritimovirus eqlidense* | WEqMV | EF608612 |
| | Yellow oat-grass mosaic virus | *Tritimovirus triseti* | YOgMV | KF985446 |
| **Hypovirus** | Hypovirus cacaofunestae | | HVc | BK062942 |
| | Monilinia fructicola hypovirus 2 | | MoFHV2 | ON038401 |
| | Cryphonectria hypovirus 1 | *Alphahypovirus cryphonectriae* | CHV1 | NC_001492 |
| | BlackFly associated hypovirus | | BFAHV | PV236822 |
| | Mycosphaerella hypovirus A | | MpHV1 | MK231018 |

***GenBase accession**
