## Supplemental Figures for "Discovery of novel members of the *Potyviridae* family reveals expanded diversity, a broad host range, and evidence of fungal and oomycete infections"

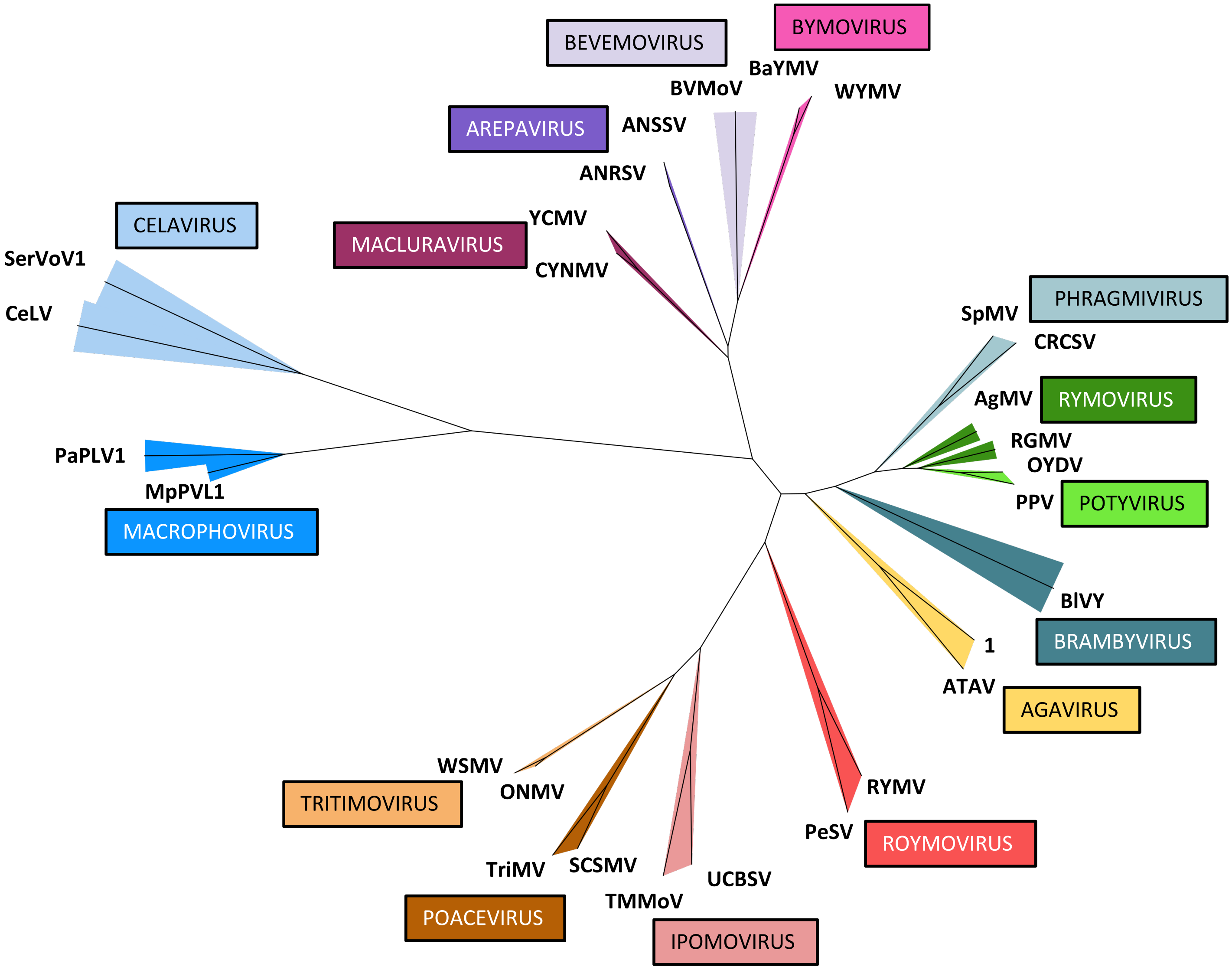


Supplementary Figure 1

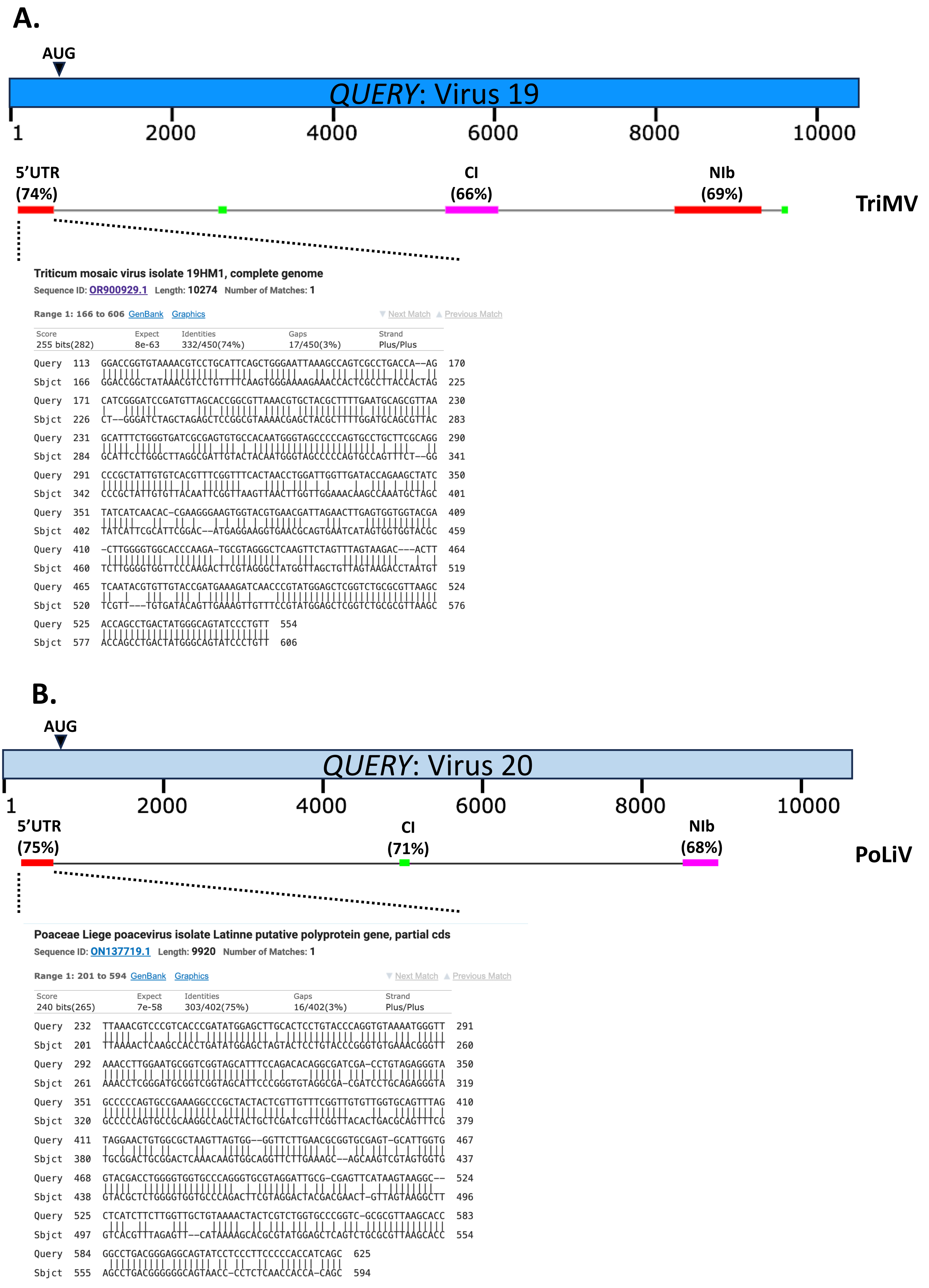


**Supplementary Figure 2**

**A.**

#### Virus 36 (*Potyvirus* genus)

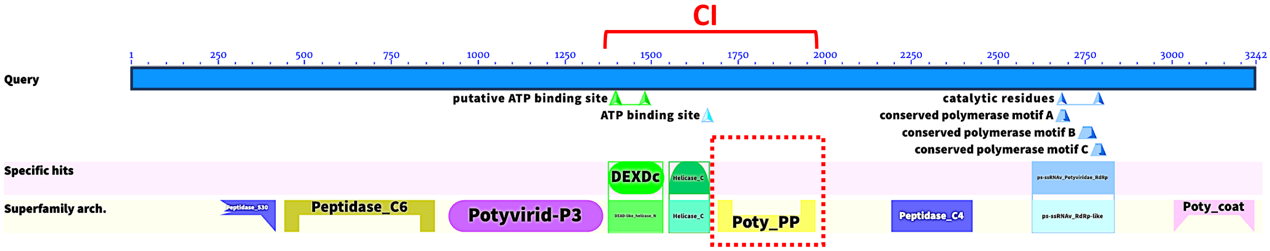


| Name | Accession | Description | Interval | E-value |
|---|---|---|---|---|
| + ps-ssRNAv_Potyviridae_RdRp | cd23175 | catalytic core domain of RNA-dependent RNA polymerase (RdRp) in the family Potyviridae of positive-sense single-stranded RNA [(+)ssRNA] viruses | 2598-2833 | $1.26 \times 10^{-172}$ |
| + Peptidase_C6 | pfam00851 | Helper component proteinase | 442-875 | $6.64 \times 10^{-118}$ |
| + Poty_coat | pfam00767 | Potyvirus coat protein | 3006-3237 | $2.60 \times 10^{-105}$ |
| + Potyvirid-P3 | pfam13608 | Protein P3 of Potyviral polyprotein | 915-1357 | $1.11 \times 10^{-73}$ |
| + Poty_PP | pfam08440 | Potyviridae polyprotein | 1691-1972 | $5.99 \times 10^{-64}$ |
| + Peptidase_C4 | pfam00863 | Peptidase family C4 | 2192-2425 | $2.12 \times 10^{-48}$ |
| + Peptidase_S30 | pfam01577 | Potyvirus P1 protease | 256-416 | $8.11 \times 10^{-28}$ |
| + DEXDc | smart00487 | DEAD-like helicases superfamily | 1376-1532 | $8.96 \times 10^{-22}$ |
| + Helicase_C | pfam00271 | Helicase conserved C-terminal domain | 1551-1666 | $3.48 \times 10^{-13}$ |

**B.**

#### Virus 10 (*Macrophovirus* genus)

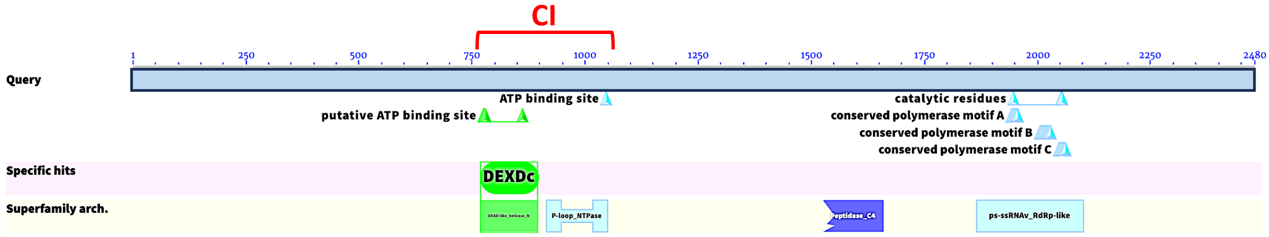


| Name | Accession | Description | Interval | E-value |
|---|---|---|---|---|
| + ps-ssRNAv_Potyviridae_RdRp | cd23175 | catalytic core domain of RNA-dependent RNA polymerase (RdRp) in the family Potyviridae of positive-sense single-stranded RNA [(+)ssRNA] viruses | 1866-2101 | $4.00 \times 10^{-57}$ |
| + SF2_C_viral | cd18806 | C-terminal helicase domain of viral helicase | 915-1049 | $1.65 \times 10^{-9}$ |
| + DEXDc | smart00487 | DEAD-like helicases superfamily | 769-894 | $2.79 \times 10^{-8}$ |
| + Peptidase_C4 | pfam00863 | Peptidase family C4 | 1526-1659 | $1.89 \times 10^{-7}$ |

**Supplementary Figure 3**

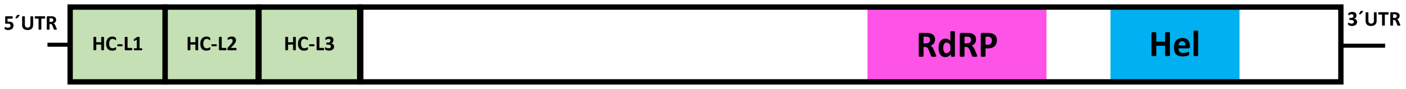


| | | | |
|---|---|---|---|
| HVc | VG \| S | VG \| G | VG \| G |
| MpHV1 | VG \| S | LG \| A | VG \| M |
| MoFHV2 | XX | VG \| G | VG \| G |
| BFAHV | XX | VG \| G | VG \| G |

**Supplementary Figure 4**

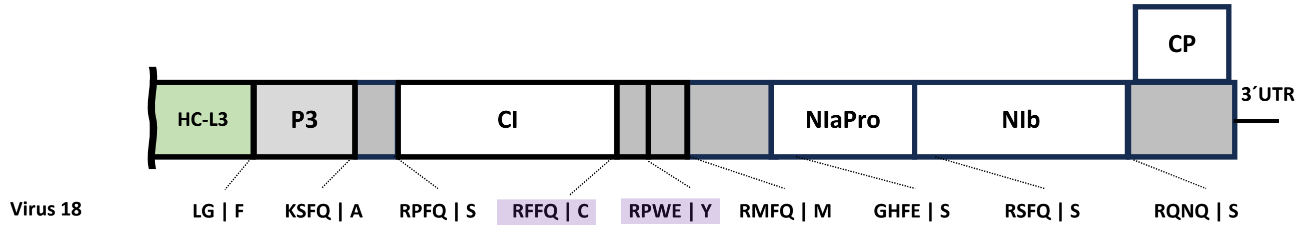


**Supplementary Figure 5**

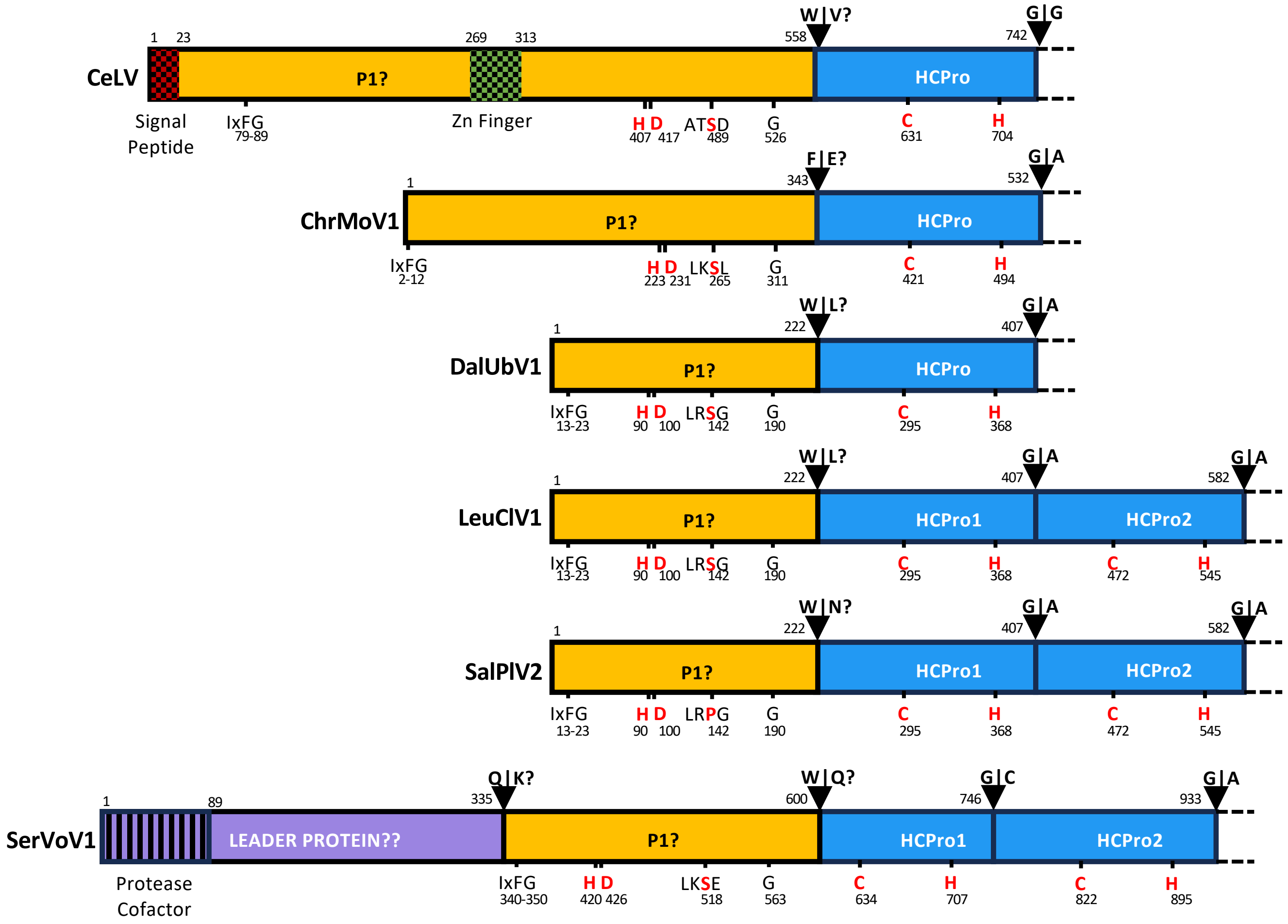


### Supplementary Figure 6
